## Supplementary Materials for "Deciphering circulating tumor cells binding in a microfluidic system thanks to a parameterized mathematical model"

June 26, 2024

### A Numerical modeling of the fluid

#### A.1 Validation of the hydrodynamic entrance length $\ell$

Let us define the domain $\Omega := (0, L) \times (0, h) \times (0, l)$, $\partial\Omega$ its boundary and $\partial\Omega_{in} := \{0\} \times (0, h) \times (0, l)$ the boundary corresponding to the channel inlet. We would like to numerically establish the distance $\ell$ (along the $x-$axis) at which a uniform velocity profile from $\partial\Omega_{in}$ fully develops into a Poiseuille profile. To do so, we compare the solution of the two following problems: (i) a Poiseuille profile is already developed over the entire domain $\Omega$, (ii) boundary layer equations are used to take into account the entrance effect near the inlet. Let $u_P$ be the fluid velocity anywhere in $\Omega$ according to the Poiseuille equations

$$\begin{cases} \rho \dfrac{\partial u_P}{\partial t} - \mu \Delta u_P = G(1 + \xi_f \cos(\omega_f t + \varphi)) & t > 0,\ \Omega, \\ u_P(t, \cdot) = 0 & t > 0,\ \partial\Omega, \\ u_P(0, \cdot) = 0 & \Omega, \end{cases} \tag{1}$$

and $u_B$ be the fluid velocity anywhere in $\Omega$ according to the boundary layer equations

$$\begin{cases} \rho \left( \dfrac{\partial u_B}{\partial t} + u_B \nabla \cdot u_B \right) - \mu \Delta u_B = G(1 + \xi_f \cos(\omega_f t + \varphi)) & t > 0,\ \Omega, \\ u_B(t, \cdot) = 0 & t > 0,\ \partial\Omega \backslash \partial\Omega_{in}, \\ u_B(t, \cdot) = \max\limits_{t>0} u_P & t > 0,\ \partial\Omega_{in}, \\ u_B(0, \cdot) = 0 & \Omega. \end{cases} \tag{2}$$

[1]Inria, Univ. Bordeaux, CNRS, Bordeaux INP, IMB, UMR 5251, F-33400 Talence, France
[2]INSERM UMR_S 1109, Univ. Strasbourg, FMTS, Équipe labellisée Ligue Contre le Cancer, F-67000 Strasbourg, France.
[1]Inria, Univ. Bordeaux, CNRS, Bordeaux INP, IMB, UMR 5251, F-33400 Talence, France
[2]INSERM UMR_S 1109, Univ. Strasbourg, FMTS, Équipe labellisée Ligue Contre le Cancer, F-67000 Strasbourg, France.

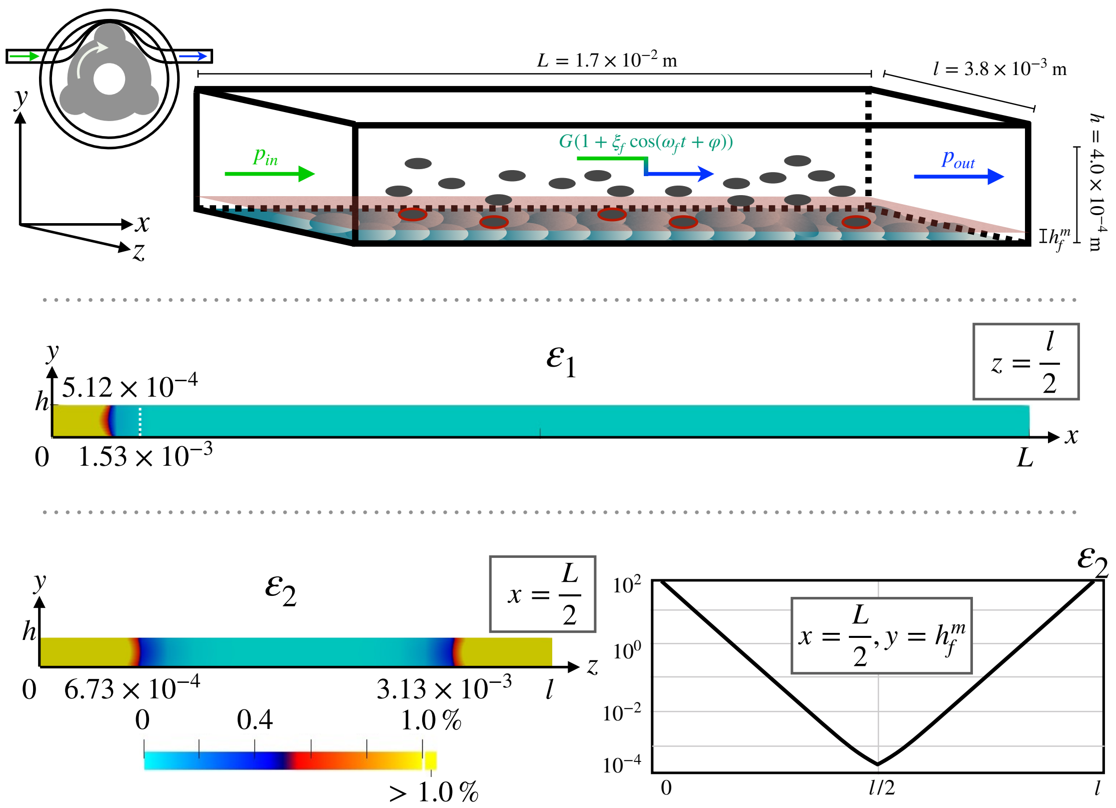


Figure 1: Top: Experimental setup. Middle: Relative error $\varepsilon_1$ on the cross-section $(0, L) \times (0, h) \times \{\frac{l}{2}\}$. For $x \geq 5.12 \times 10^{-4}$ m we have, $\varepsilon_1(\cdot) \leq 1\%$. For reference, the entry length $\ell = 1.53 \times 10^{-3}$ m given by [47] is also displayed. Bottom-Left: Relative error $\varepsilon_2$ on the cross-section $\{\frac{L}{2}\} \times (0, h) \times (0, l)$. For $z \in [6.73 \times 10^{-4}\ \text{m}, 3.13 \times 10^{-3}\ \text{m}]$ we have, $\varepsilon_2(\cdot) \leq 1\%$. Bottom-Right: Relative error $\varepsilon_2$ on the cross-section $\{\frac{L}{2}\} \times \{h_f^m\} \times (0, l)$

The third equation of Problem (2) corresponds to a uniform velocity profile at the inlet of the channel, which is here equal to the maximum velocity of the fluid with a fully developed Poiseuille profile. This choice was made to be consistent with the steady state hypothesis. Problems 1 and 2 are numerically solved using `FreeFEM++` [53]. The domain is discretized on a $120 \times 10 \times 20$ mesh, using P1 Lagrange elements. The timestep $\Delta t$ is taken to be 0.5 s and the total time of the simulation is 15 s.

For the parameters, we work with the *worst case scenario*, *e.g.* the highest Reynolds number possible, with $Re = 3.75$. We recall that $L = 1.70 \times 10^{-2}$ m, $h = 4.00 \times 10^{-4}$ m, $l = 3.80 \times 10^{-3}$ m, $\rho = 1.00 \times 10^{3}$ kg.m$^{-3}$, $\mu = 7.20 \times 10^{-4}$ Pa.s, $G = 201.32$ Pa.m$^{-1}$, $\omega_f = 4.88 \times 10^{1}$ rad.s$^{-1}$, and $\varphi = 0$ rad, see Subsections 2.2, 3.2 and 4.2 for details. By convention, a flow can be considered fully developed when its velocity profile matches the asymptotic one with an error margin of less than a percent. Assuming $u_B(t, \cdot) \neq 0$, for all $t > 0$, the relative error in percent is computed as follows

$$\varepsilon_1(\cdot) := \max_{t>0} \left( \frac{|u_B(t,\cdot) - u_P(t,\cdot)|}{|u_B(t,\cdot)|} \right) \times 100, \quad \Omega.$$

Figure 1-Middle shows the relative error $\varepsilon_1$ over the longitudinal cross-section $(0, L) \times (0, h) \times \{\frac{l}{2}\}$. One may remark the hydrodynamic entrance length $\ell$ – displayed along the cross-section – seemingly over-predicts the distance at which $\varepsilon_1$ drops below one percent. The numerical entrance length at which $\varepsilon_1 \leq 1\%$ is given by $\ell_{num} = 5.12 \times 10^{-4}$ m. Numerical tests including spatial and temporal convergence were performed to validate the value of $\ell_{num}$.

From Subsection 3.2, we know the data was obtained $\frac{L}{2} = 8.50 \times 10^{-3}$ m away from the inlet, which is a full order of magnitude above the upper bound $\ell$ of the hydrodynamic entrance length. Under those circumstances, the velocity profile of $u_B$ closely matches the velocity profile of $u_P$ and is considered as a fully developed Poiseuille flow.

**Remark A.1.** *The value* $\ell = 1.53 \times 10^{-3}$ m *given in Subsection 5.2 and coming from [47] is superior to our numerical evaluation of the hydrodynamic entrance length* $\ell_{num}$. *Furthermore, additional hydrodynamic entrance length comparisons were performed with other formulas available in the literature (see [47], Table 1), and all of them were found to overpredict* $\ell_{num}$. *This difference is explained by the relation between the velocity of the fluid far from the inlet and its velocity at the inlet boundary. As shown in [47], their study was performed for a ratio whose maximum is between* 1.5 *and* 2, *while in our study we set this maximum at* 1. *Therefore, their entry velocity is much higher than the asymptotic velocity of the fluid, which is reflected in the reported value of* $\ell$.

### A.2 Validation of the fluid expression

In Subsection A.1, we have shown the device was *long enough* for a Poiseuille flow to become fully developed and therefore independent on the $x-$axis. While this reduces Problem (1) into a 2D one, we can simplify it even further by showing the channel is *wide enough* that we can neglect a change in the $z-$axis as well. We recall that aspect ratio $AR = \frac{h}{l}$ equals to $1.05 \times 10^{-1}$, which tells us the microfluidic device is roughly 10 times wider than it is tall. To show that 1D reduced model (in $y$-axis direction) is reasonable, we compare the fluid velocity $u_P$ to the expression $u_f$, which must verify

$$\begin{cases} \rho \dfrac{\partial u_f}{\partial t} - \mu \dfrac{\partial^2 u_f}{\partial y^2} = G(1 + \xi_f \cos(\omega_f t + \varphi)) & t > 0, \ (0, h), \\ u_f(t, \cdot) = 0 & t > 0, \ (0, h), \\ u_f(0, \cdot) = 0 & (0, h). \end{cases} \tag{3}$$

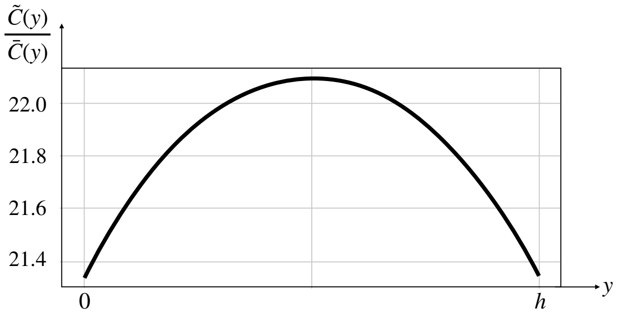


Figure 2: Quotient $\dfrac{\overline{C}(y)}{\widetilde{C}(y)}$ for $y \in (0,h)$.

Using the same parameter values and mesh as in Subsection A.1, we solve Problem (3) with `FreeFEM++`. The same error threshold of one percent is taken, with

$$\varepsilon_2(\cdot) := \max_{t>0}\left(\frac{|u_P(t,\cdot) - u_f(t,\cdot)|}{|u_P(t,\cdot)|}\right) \times 100, \quad (0,h)\times(0,l).$$

Figure 1-Bottom-Left gives $\varepsilon_2$ on the lateral cross-section $\{\frac{L}{2}\}\times(0,h)\times(0,l)$ corresponding to the section from which the data were obtained. For $z \in [6.46\times10^{-4}\ \text{m}, 3.15\times10^{-3}\ \text{m}]$ we have, $\varepsilon_2(\cdot) \leq 1\%$. Figure 1-Bottom-Right shows the relative error $\varepsilon_2$ at the camera's focal plane, using the value of $h_f^m$ found in Subsection 4.2. As expected from the observation made in Figure 1-Bottom-Left, the distance between the two inner ticks does not increase nor decrease in any significant way. The sub$-1\%$ relative error area thus spans most of the microfluidic device, with $\{\frac{L}{2}\}\times(0,h)\times(6.46\times10^{-4}, 3.15\times10^{-3})$. Upon closer inspection of the data provided in [14, 42], we know the data was obtained at the middle of the device length wise, but also width wise, as the lateral sides of the device are not visible in the videos. Given the width of the sub$-1\%$ area, $u_P$ can therefore be represented by $u_f$.

### A.3 Validation of the parabolic shape for the velocity profile

For a purely oscillating pressure gradient, a parabolic velocity profile can be considered for a **Wo** up to 1, after which the profile rapidly evolves into a *plug*-like shape [48]. Counting the number of oscillations in Figure 3, it can be seen that $\omega_f$ depends on the cohort. The highest value is reached at the third cohort and is close to 50 $\text{s}^{-1}$ ($\sim$12 oscillations on 1.5 s gives $\sim 12/1.5\times2\pi \sim$50 $\text{s}^{-1}$), resulting in a **Wo** around 1.6. However, it is shown in [49] that a parabolic profile can still emerge for moderately low values of **Wo** ($\approx 2$) when

$$|\text{Re}(u_f(t,y))| \gg |\text{Im}(u_f(t,y))| \qquad t>0, \quad y\in(0,h).$$

Using the superposition principle, one can rewrite $u_f$ as a sum of its constant and oscillating part, that we respectively denote by $\overline{u}_f(t,y)$ and $\widetilde{u}_f(t,y)$. We now have to check that

$$|\text{Re}(\widetilde{u}_f(t,y))| + |\text{Re}(\overline{u}_f(t,y))| \gg |\text{Im}(\widetilde{u}_f(t,y))| + |\text{Im}(\overline{u}_f(t,y))| \qquad t>0, \quad y\in(0,h)\,. \tag{4}$$

From Equation (2), one has $|\text{Re}(\overline{u}_f(t,y))| = \overline{C}(y) > 0$ and $|\text{Im}(\overline{u}_f(t,y))| = 0$, so that the condition rewrites

$$|\text{Re}(\widetilde{u}_f(t,y))| + \overline{C}(y) \gg |\text{Im}(\widetilde{u}_f(t,y))| \qquad t>0, \quad y\in(0,h)\,,$$

which is true for all $t$ if $\overline{C}(y)$ is taken large enough. In the most pessimistic case, we have $|\mathrm{Re}(\widetilde{u}_f(t,y))| = 0$ and $\max_{t>0} |\mathrm{Im}(\widetilde{u}_f(t,y))| = \widetilde{C}(y) > 0$. Therefore, it is enough to show that

$$\frac{\overline{C}(y)}{\widetilde{C}(y)} \gg 1.$$

Figure 2 shows the value of $\frac{\overline{C}(y)}{\widetilde{C}(y)}$ for $y \in (0, h)$. It can be deduced that Condition (4) is always verified. This means that for $u_f = \overline{u}_f + \widetilde{u}_f$, the constant component is large enough to ensure that the magnitude of oscillations does not disrupt the parabolic shape of the velocity profile. This therefore validates our hypothesis.
